# Chaotic neural dynamics facilitate probabilistic computations through sampling

**DOI:** 10.1101/2023.05.04.539470

**Authors:** Yu Terada, Taro Toyoizumi

## Abstract

Cortical neurons exhibit highly variable responses over trials and time. Theoretical works posit that this variability arises potentially from chaotic network dynamics of recurrently connected neurons. Here we demonstrate that chaotic neural dynamics, formed through synaptic learning, allow networks to perform sensory cue integration in a sampling-based implementation. We show that the emergent chaotic dynamics provide neural substrates for generating samples not only of a static variable but also of a dynamical trajectory, where generic recurrent networks acquire these abilities with a biologically-plausible learning rule through trial and error. Furthermore, the networks generalize their experience in the stimulus-evoked samples to the inference without partial or all sensory information, which suggests a computational role of spontaneous activity as a representation of the priors as well as a tractable biological computation for marginal distributions. These findings suggest that chaotic neural dynamics may serve for the brain function as a Bayesian generative model.

---

Humans and other animals face environments inherently associated with uncertainty, necessitating the handling and integration of uncertain information for survival. A wide spectrum of experimental studies has shown that the brain can perform nearly optimal Bayesian computation [1–6]. While some computational models [2, 7] assume that neurons encode statistics of the underlying probability distribution, others suggest that neurons encode Monte Carlo samples drawn from the distributions [8–15]. For the latter models, variability in neural activity is an essential element for probabilistic information representation.

Consistently, recent experiments have recorded a large number of neurons simultaneously and revealed that irregular patterns of neural activity [16, 17] underlie information processing in the brain. Such variability is generated spontaneously even in the absence of explicit changes in sensory input [18–20]. At a macroscopic scale, functional magnetic resonance imaging observations reveal specific patterns of intrinsic variability during the resting state. These patterns, known as default mode networks [21], exhibit structures that reflect experience and knowledge [22]. At a microscopic scale, neural avalanches, in which neurons exhibit events with strong synchrony and burst-type activity with power-law distributions of sizes and lifetimes, are observed during spontaneous neural activity [23]. Thus, irregular spontaneous neural activity ubiquitously emerges across various spatiotemporal scales.

The biological source of neural variability is currently under debate [24–26]. Neural variability has sometimes been modeled by different types of stochastic noise in neural dynamics [27–29], such as input noises to neurons [14] or stochastic spiking due to spike-threshold noise in point-process-type models [30]. Another factor of neural variability arises from vesicular transmission by synapses [31], which is modeled by vesicular release probability. These works assume the existence of a random number generator separately from the modeled neural circuits. In contrast to these stochastic models, our study focuses on models that explain neural variability in deterministic systems [32–38]. Strong and heterogeneous synaptic connections with the overall balance between excitatory and inhibitory drives can endow even deterministic neural networks with the ability to generate high-dimensional variability by chaos. Some experimental observations support this hypothesis [25]. This line of research has also triggered theoretical works elucidating the computational advantages of neural dynamics at the edge of chaos [39–42]. Further, chaotic network states are shown to be a suitable initial condition that enhances learning efficiency [43–46]. However, these neural networks do not typically exhibit chaos after learning, and their chaoticity has not been utilized explicitly for computation in trained networks. More recently, chaotic dynamics are shown to permit multiple time scales even close to marginally stable systems, which is advantageous for enabling long short-term memory without finely tuning parameters [38, 47].

Previous Bayesian computation models of the brain [7–15] assume externally injected noise as the source of variability. Hence, whether chaotic variability is compatible with biological Bayesian computation is unclear, especially because the chaotic property that expands the effect of perturbation in time is apparently contradictory to accurate computation. In addition, these stochastic models often rely on a parametric family to describe target probability distributions, which requires hand-crafted designs in advance. In biological setups, considering generic network architecture at the early stage of learning would be more plausible. While non-parametric methods to learn a target probability distribution are popular in machine learning [48–51], it is not trivial how the brain can implement these computations in a biologically plausible manner.

Here, we consider a non-parametric model to generate samples from the Bayesian posterior distribution. The sampling is achieved by utilizing chaotic network dynamics of recurrently connected deterministic neurons. During training, we use the biologically plausible node perturbation learning rule [52–54], which is represented as a “three-factor learning rule.” Three-factor learning rules are promising candidates for enabling flexible adaptation for biological neural networks in a broad range of tasks [55–58] as they can be implemented with local computation, namely, by a Hebbian learning rule modulated by a global signal [59–61]. We explore the efficiency of a three-factor learning rule.

We consider representative examples of Bayesian tasks to demonstrate that chaotic neural dynamics serve as the substrate for representing posterior probability distributions. Specifically, we use paradigmatic cognitive tasks for integrating information from multiple sources similar to human and animal tasks, known as cue integration tasks [62, 63]. We demonstrate that, after training using a local learning rule, chaotic recurrent neural networks learn to sample hidden static states or dynamic trajectories of a hidden variable in a near-optimal manner. We discuss the implications of using chaos for probabilistic computations in the brain.

## Results

### Neural sampling through chaotic dynamics

We developed a rate-based recurrent network model that uses their intrinsic dynamics to draw samples of the hidden variable of interest from an estimated Bayes posterior distribution. These neurons are governed by discrete-time dynamics receiving inputs from their recurrent synapses and sensory neurons (see below and Methods). A simple cue-integration task is described in Fig. 1 a. In this task, recurrent networks infer the probability distribution of the hidden variable *θ* of interest. For example, *θ* may describe the angular position of an object. For simplicity, we assume that the angular position can take one among *n* possible directions (A biologically plausible extension will be discussed in Discussion). A recurrent neural network receives two kinds of input vectors **x**^*α*^, which represent the firing patterns of two sets of sensory neurons *α* = *A, B*. For example, the sets A and B may represent ensembles of auditory and visual sensory neurons, respectively, that encode the direction *θ* of the object. The lengths of these input vectors are also assumed *n*, and each element *k* takes either the active or inactive state, that is, 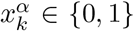. At the beginning of each trial, the value of *θ* is randomly drawn from a uniform distribution unless stated otherwise (non-uniform cases will be shown later), and the input elements are generated independently according to the conditional probability 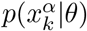 for an element *k* = 1, 2, …, *n* and population *α*. This conditional probability specifies the tuning curve of neurons, characterizing the sensory neurons’ activation probability given the angular position. It peaks at *k* = *θ* and decreases (slowly for *α* = *A* and rapidly for *α* = *B*) as *k* deviates from the peak (an example is shown in Fig. 1 b; see also Methods). Note that the network receives the information of *θ* only through noisy firing patterns **x**^*α*^ of the sensory neurons. Within each trial of its length *T* = 200 time steps, the values of *θ* and **x**^*α*^ are fixed in time. The discrete-time dynamics of the *N* recurrently connected neurons at time *t* (*t* = 1, 2, …, *T*) are described by **h**(*t*) = *Jϕ* (**h**(*t −* 1)) + Σ*α*=*A,B K*^*α*^**x**^*α*^ + **c**, where **h** is a *N* -dimensional vector of recurrent neural network activity, *J* is a *N × N* matrix of recurrent synaptic weights, *K*_*α*_ (*α* = *A, B*) are the *N × n* synaptic weight matrices from input *α*, **c** is a *N* -dimensional vector of baseline input parameters, and *ϕ*(*·*) = tanh(*·*) is an activation function (see also Methods). The entries of *J* are initially drawn identically and independently from the Gaussian distribution with zero mean and standard deviation 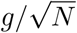, parameterized by the gain parameter *g*. Note that both *J* and **c** change during learning. The entries of *K*_*α*_ are drawn from the standard normal distribution, and they are fixed during the entire simulation. Finally, an *n*-dimensional vector **y**(*t*) represents the binary (0 or 1) activity of output neurons at time *t*. These neurons receive inputs **z**(*t*) = *Wϕ*(**x**(*t*)) + **b**, where *W* denotes *n × N* matrix of readout weights and **b** is an *n*-dimensional vector of bias parameters, and only the output neuron that receives the maximum input is active while the remaining are inactive, which could be implemented by the winner-take-all (WTA) mechanism [64]; the active output neuron at time *t* represents a sample of the hidden variable, namely, 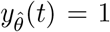 and 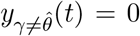 with 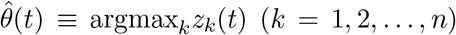. Our goal is to train the network so that the network-generated histogram of 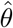 approximates well the Bayesian posterior *p*(*θ*|**x**^*A*^, **x**^*B*^) given the current sensory input pattern.

**Fig. 1:**
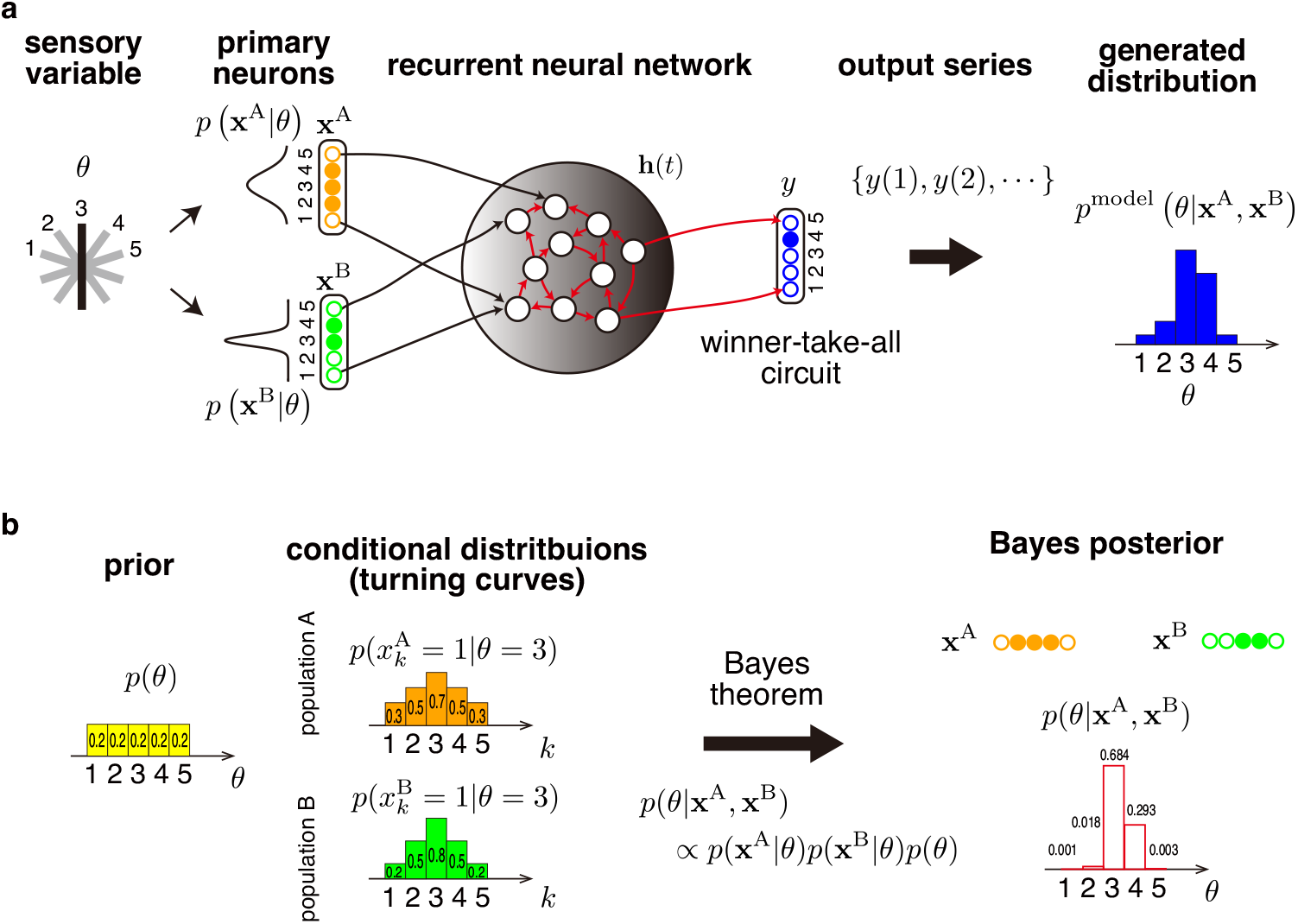
**a**, Schematics of the cue integration task. The recurrent neural network receives inputs **x**^*A*^ and **x**^*B*^ as sensory inputs, whose firing patterns are determined stochastically based on their tuning curves and a hidden variable *θ* of interest. The network learns to infer the probability distribution of *θ* given these inputs and generates samples through the output activity. The network’s output histogram composes an estimate of the posterior probability. During the learning period, the bias parameters as well as the internal and readout connections in red are modified. **b**, The Bayes optimal posteriors in the cue integration task. The prior of *θ* (the uniform distribution here) is shown with the yellow histogram on the left. The conditional distributions of the two input vectors are specified by their tuning curve properties. The tuning curves describe the probability that each input neuron is active given *θ*; they are depicted using the orange and green histograms in the middle for *θ* = 3. The Bayes posterior distribution given these input patterns **x**^*A*^, **x**^*B*^ is specified by the Bayes theorem and indicated by the red blank histogram on the right.

We adopted a local learning rule based on the node perturbation method [52–54] to update synaptic weights and bias parameters, which is considered more biologically plausible than machine learning algorithms that require, for example, error backpropagation (see the details in Methods). The error function for learning is defined as the square of the Hellinger distance [65], which measures the distance between the histograms of active output neurons within the time window and the Bayesian posterior. We assumed that the Hellinger distance is evaluated by an external network and transferred to the modeled network during learning. According to the node perturbation learning scheme, we updated adjustable parameters *J, W*, **c** and **b**, but set the baseline parameter **c** as zero through simulations for the static tasks. For example, the update of synapse *J*_*ij*_ is described by the product of the global signal and a Hebbian term that correlates presynaptic activities and postsynaptic perturbations (Eq. (6)). The stochastic perturbations for computing the gradient are turned off after the training, and therefore the network dynamics become completely deterministic.

While a random network does not exhibit adequate sampling before training, the trained network utilizes its irregular dynamics to represent the posterior distribution (Fig. 2). The neural variability in the recurrent network plays a crucial role in expressing the uncertainty of the hidden variable because the goal here is not only to estimate the most likely value of *θ* but also to generate samples from the posterior distribution. Moreover, the trained network succeeded in integrating uncertain information from two different sources and produced a good approximated posterior probability.

**Fig. 2:**
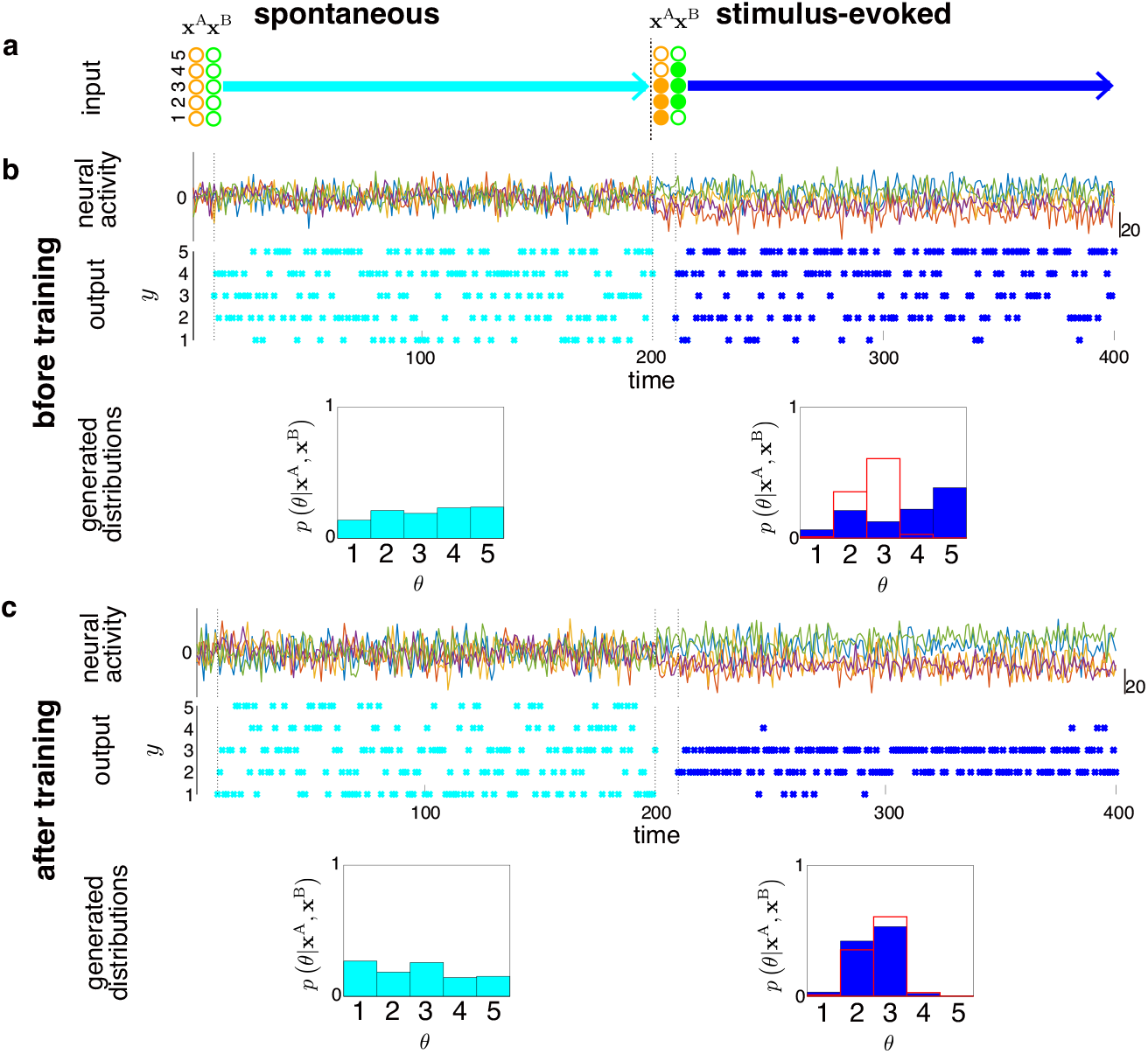
An example of neural sampling before (with random initial weights) and after training. **a**, The input patterns of the two sensory neuron populations. During the first-half period, sensory inputs are absent (all the sensory neurons are inactive). During the second-half period, the networks receive specific patterns of sensory inputs according to a hidden object direction. **b, c**, (Top) The activities of a randomly selected subset of neurons, (Middle) the activities of output neurons (blue dots indicate active neurons), and (Bottom) the resulting histograms of active output neurons (colored bars). The generated distributions for the second half with blue-filled boxes are compared with the Bayesian posterior distributions, denoted by the red blank boxes.

In Fig. 3 a, the time course of the error (the squared Hellinger distance between the generated output histogram and the Bayesian posterior) is shown for 10 randomly initialized networks during learning. The task performance depends on which sensory inputs are available. The performance using input B alone is better compared to that using input A alone. This is because the neurons in B have a narrower tuning curve, and the input B is hence more informative than input A. The best performance is achieved when both inputs are available, highlighting the ability of the network to integrate multiple cues. The resulting distributions *p*(*θ*|**x**^A^, **x**^B^) represented by the trained network for different patterns of sensory inputs, closely approximate the Bayesian posteriors over a variety of input vectors (Fig. 3 b). As expected, the output distribution approaches the Bayesian posterior as the sampling duration *T* increases (Fig. S1). Further, the adaptation of the synaptic connectivity *J* within the recurrent network reduces significantly the sensitivity of learning outcomes to the initial network configuration, compared to the so-called “reservoir scheme” that adjusts only the readout parameters *W* and **b** (see, initialization at the edge of chaos in Fig. S2 and at the chaotic regime in Fig. S3). Randomly connected recurrent neural networks with strong synaptic interactions can exhibit chaotic dynamics. To investigate the nature of network dynamics and how they are related to task performance, we monitored the largest Lyapunov exponent as a measure of chaoticity and compared it with the error function during the learning period (Fig. 3 c). The network is initially placed slightly below the edge of chaos with the synaptic gain parameter *g* = 1.05. Note that the dynamics are non-chaotic at *g* = 1.05 because the sensory inputs suppress chaos in random neural networks as demonstrated previously (e.g., [66] and Fig. S3). The result shows that, as task performance improves through learning, the largest Lyapunov exponent increases from the negative to the positive side. It indicates that chaos emerges through learning while the network develops the capacity for probabilistic computation. This result is in contrast to the findings of previous works that trained non-chaotic dynamics [43–45], and it highlights the utility of chaotic dynamics in probabilistic computation. The enhancement of chaoticity by learning was reported in associative memory [67], but our study directly links the computational ability of networks to chaos through the lens of representation of probabilistic distributions.

**Fig. 3:**
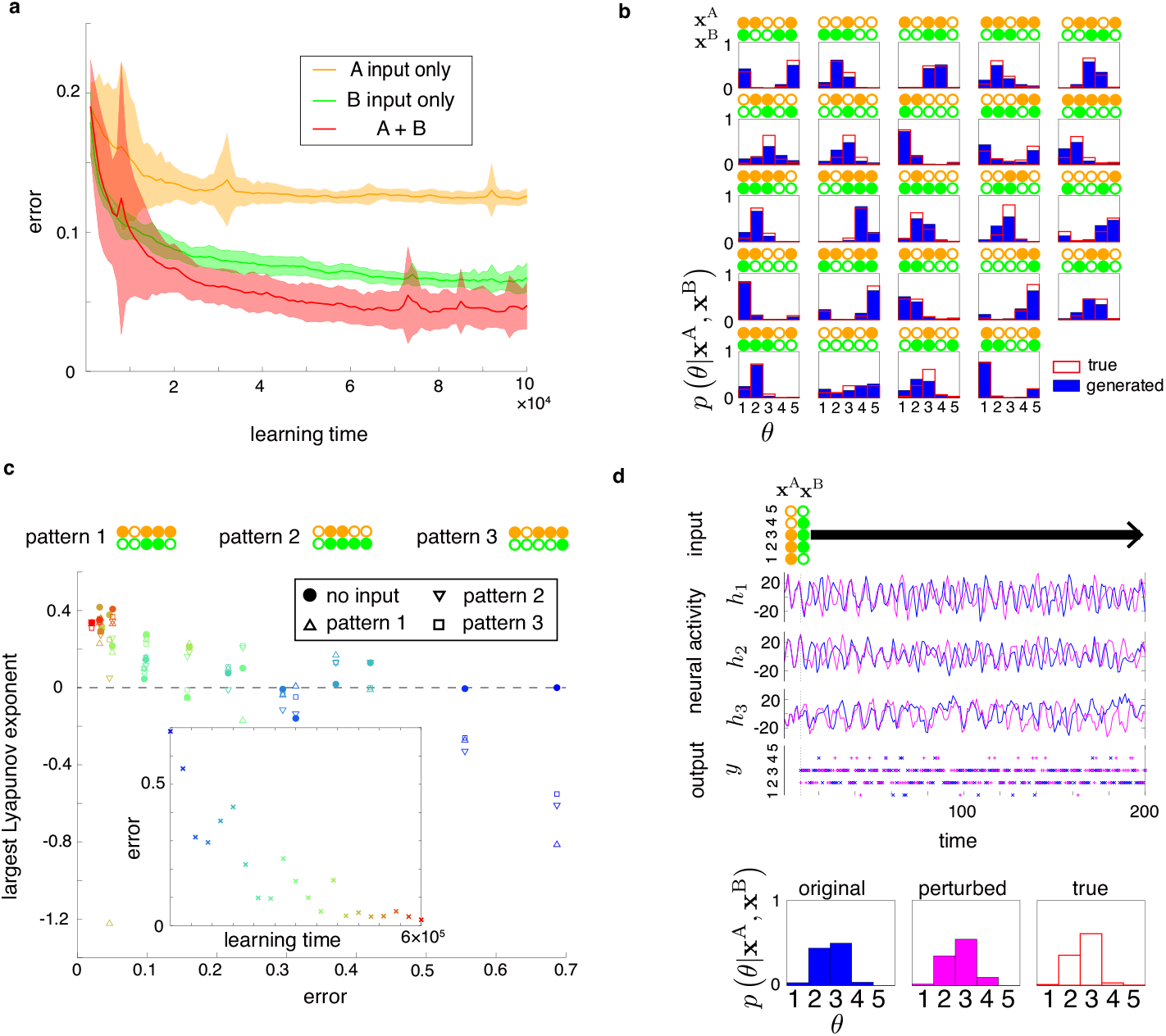
Performance and chaoticity of the recurrent neural networks in the cue integration task. **a**, Learning curves of the networks that receive inputs from either of the populations (A or B) and from both. **b**, Probability distributions provided by the trained network (the blue filled boxes) and true posteriors corresponding to the sensory inputs (red blank boxes) for multiple examples of input patterns. Input patterns A and B are shown above the distributions. **c**, Largest Lyapunov exponent versus error value for networks during training for different patterns of inputs. The inset shows the errors versus learning time. The markers are colored from blue to red depending on the elapsed time. **d**, Top: Input, dynamics of selected neurons, and output in the trained networks with two similar initial conditions, where the initial state of the magenta trajectory is slightly perturbed from that of the blue trajectory. Bottom: Estimated posterior distributions generated by the two networks and the true posterior.

Chaotic neural dynamics render the networks to be highly sensitive to a perturbation, which might appear incompatible with accurate computation. Indeed, when the initial activity of neurons **h**(0) is slightly perturbed in the trained network, its subsequent trajectory changes drastically (Fig. 3 d). Nonetheless, the output distributions in response to various sensory inputs are robust to the perturbation (see also Fig. S4). Hence, the high sensitivity to the initial condition does not compromise the reliability of the probabilistic computation. Furthermore, we demonstrate robust performance against perturbations in recurrent synaptic weights *J* (Fig. S5). As we employ the gradient method in learning, we implicitly assume that a perturbation in synaptic weights *J* does not generate divergence; it is confirmed in Fig. S5. We believe that this is a possible reason why the brain can perform reliable computation using highly variable neural dynamics.

### Computational role of spontaneous and evoked neural activities

As shown above, the recurrent networks draw output samples from distinct sets during spontaneous and evoked activities. The spontaneous activity of neurons may reflect the prior distribution over possible sensory inputs [5, 9]. Here, we use our model to test the hypothesis that the learning process through stimulus-evoked samples can construct priors. In Fig. 2 c, we assumed the uniform prior distribution over *θ* and obtained a near uniform output distribution during the spontaneous activity. However, the output distribution during spontaneous activity looked similar to the uniform prior even before the learning (Fig. 2 b). Moreover, the learning process explicitly supervised the prior distribution by giving the networks training samples, including input sets without sensory stimulus. To address the problem of whether the network learns to code the prior distribution *p*(*θ*) during spontaneous activity from the learning only with stimulus-evoked samples, we trained networks using two different prior distributions (Fig. 4 a, b). The results show that the histogram of output during spontaneous activity matches the prior distribution *p*(*θ*) well. The learning curves for the errors for stimulus-evoked and spontaneous patterns indicate that the local learning sculpts chaotic dynamics not only for the posteriors but also for the prior only relying on stimulus-evoked samples (Fig. 4 c).

**Fig. 4:**
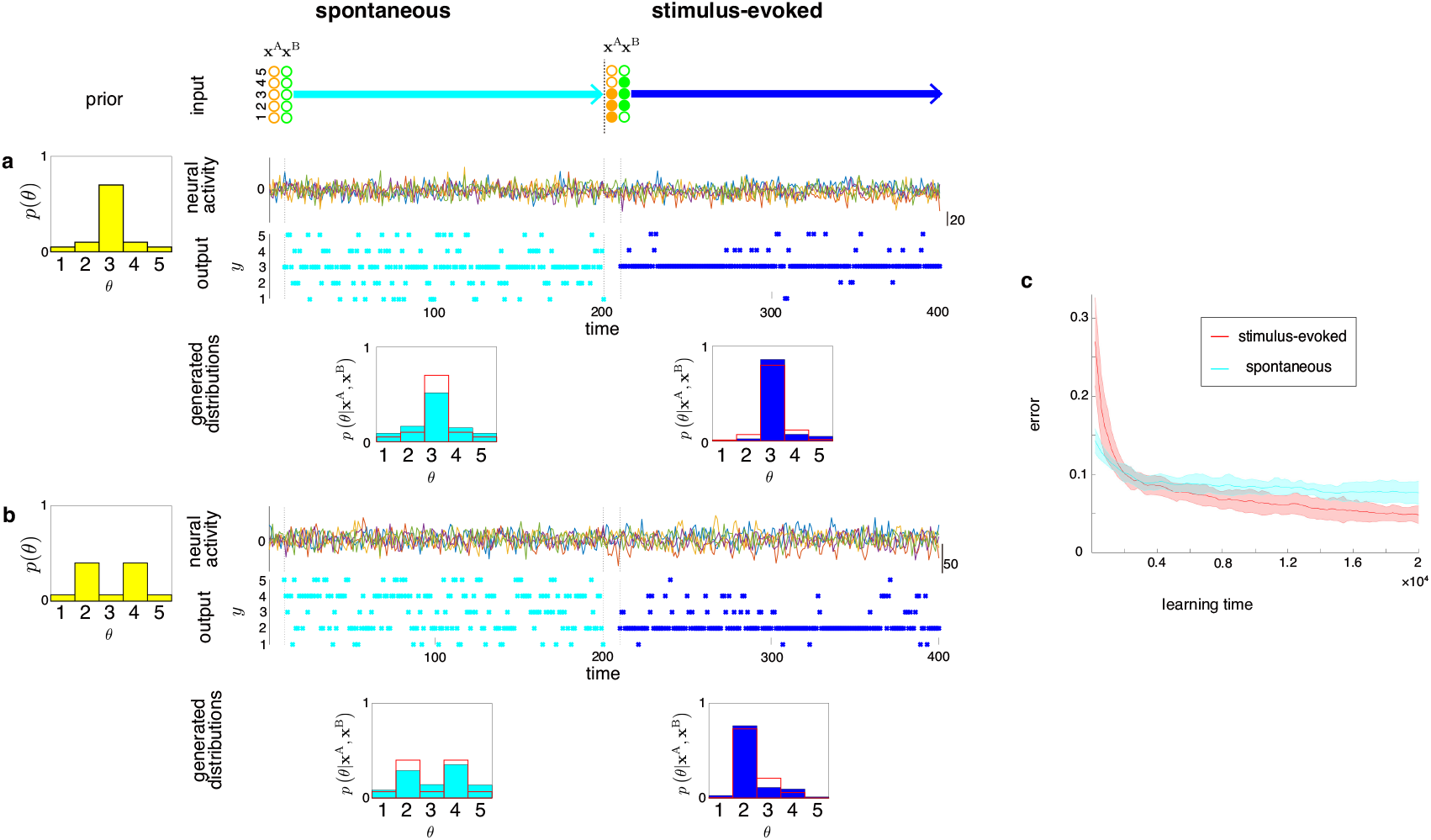
**a, b**: Neural sampling after learning with different priors. Sensory inputs are absent during the first-half period while a specific pattern of sensory inputs is injected during the second-half period. Two example priors, for **a** and **b**, are represented by the yellow histograms on the left, respectively, which are distinct from the uniform distribution used in Fig. 2. The generated distributions during the two periods are shown below the neural activity traces and the outputs on the right. The conventions are as in Fig. 2. **c**: Learning curves for training errors with stimulus-evoked cases (red curve) and generalization errors where sensory inputs are absent (cyan curve). The true prior is the same as **a**.

We showed above that the network output is distributed approximately representing the prior distribution when both sensory inputs are absent. Note that the prior distribution is written by averaging the posterior distribution with respect to the two unobserved sensory inputs, 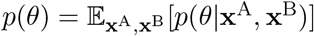, where 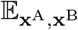 represents the expectation over **x**^A^ and **x**^B^. A natural extension of the above result is the case, where only one of the sensory inputs is observed. For example, if only **x**^A^ is observed, the output distribution might follow the conditional average of the posterior of the form 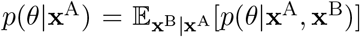, where 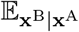 is the conditional average over **x**^B^ given **x**^A^. Our simulations demonstrate that this property holds with high accuracy. Figure 5 shows the result in two cases where the trained networks receive inputs either from population A or B. We note that, as in Fig. 4, the networks were trained using only stimulus-evoked samples. Namely, the network receives inputs from at least one active neuron of each sensory population. This situation can correspond to a condition in which an animal receives partial information (for example, only auditory or visual inputs) and generalize its experience to this novel situation with more uncertainty. Conditionally averaging the posterior distribution in a direct manner can be computationally expensive because it involves the sum over potential unobserved input candidates, whose complexity grows exponentially with the number of unobserved variables. Moreover, the computation must revise when the observed input changes. Through chaotic sampling, the networks succeed in approximating the conditionally averaged posteriors circumventing the evaluation of the sum over unobserved input candidates. This implementation constitutes a biologically suitable strategy to implement probabilistic computation with limited computational resources.

**Fig. 5:**
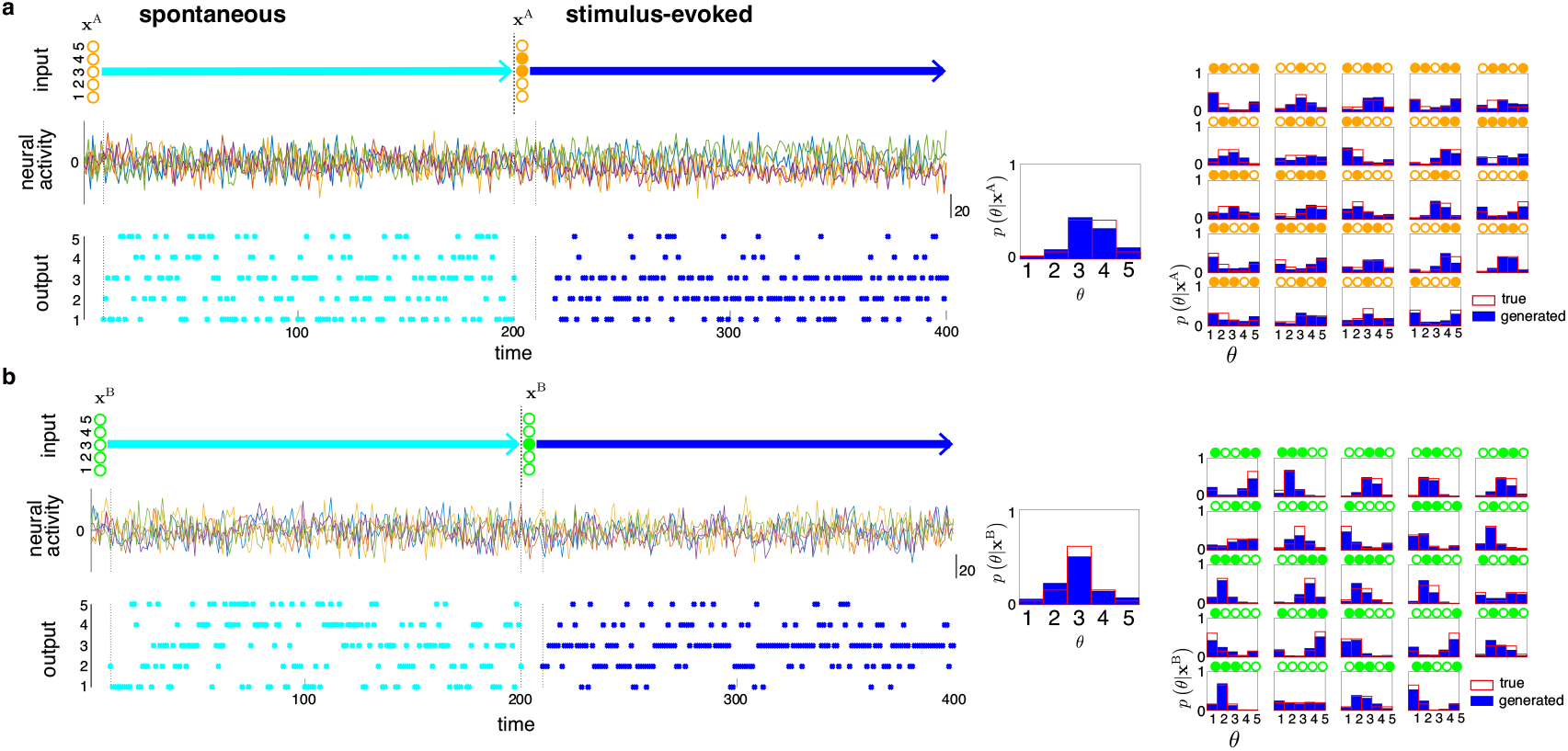
Inference by the trained networks under a single cue. Left: Neural sampling for two cases given a single cue from A or B neurons in the trained networks. Sensory inputs are absent during the first-half period, and the networks receive the inputs only from one of the two populations (a: A, b: B population) of sensory neurons during the second-half period. The conventions are as in Fig. 2. Middle: Generated distributions and Bayes posteriors for the case in the left panel. Right: A set of examples that correspond to Fig. 3 b but with only one of the sets of sensory neurons observed.

### Dynamical probabilistic inference

Next, we illustrate another example of probabilistic tasks that recurrent neural networks can implement using their chaotic dynamics. In the previous task, the networks exhibit their ability to integrate multimodal information to infer underlying static probability distributions. We hypothesize that the recurrent network model can also learn to generate samples of dynamic trajectories in a sensory-input-dependent manner. The task we consider here is one in which the network discriminates types of inputs and, depending on them, switches the transition probabilities of the outputs (Schematics is shown in Fig. 6 a). It requires the network to generate samples depending on its current internal state. This task also includes an essence to perform Markov chain Monte Carlo computation, applicable in various computations [68, 69]. To implement this computation, recurrent neural networks are required to learn temporal statistics. Here, we consider a simple setup, in which inputs A and B are both single-bit binary inputs, and we assume *n* = 3 states of *θ* and the corresponding output. The target transition probability matrices depend on sensory inputs; they have a uniform structure without sensory inputs and heterogeneous structures with sensory inputs (see Methods). The same training scheme for the internal and readout parameters is applied again. We note that a network requires both sources of information as sensory inputs for accurate inference and, hence, this task is also a cue integration task.

**Fig. 6:**
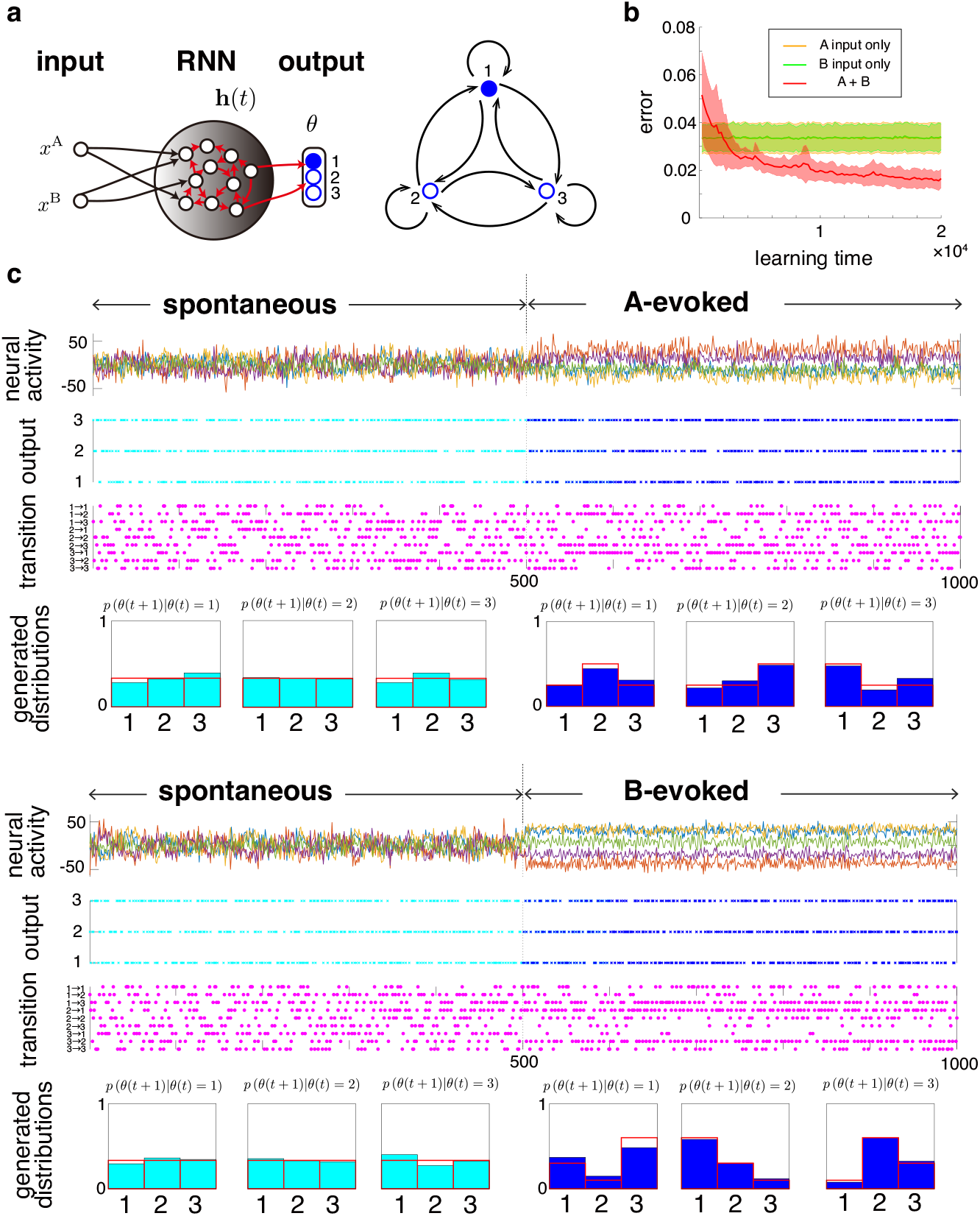
Generating sample sequences with temporal relations depending on sensory inputs. **a**, Schematics of the setting. (Left) Network setup. (Right) Markovian transitions between the output states. **b**, Learning curves using either input or both. **c**, Neural dynamics, output, transition types, and transition histograms after learning. The network receives no input in the first half and receives the sensory input (*x*^*A*^, *x*^*B*^) = (1, 0) (Top) or (0, 1) (Bottom) in the second half.

The trained recurrent networks integrate the information of the two inputs and utilize their chaotic variability to generate a dynamic output trajectory. Their learning curves are shown in Fig. 6 b. The generated output trajectories approximately follow the target transition probabilities for both types of input patterns (Fig. 6 c), while the time-averaged distributions of the outputs themselves are uninformative and almost uniform. Thus, the output dynamics driven by chaos can accurately reproduce the sequence of the target variable that follows the input-dependent stochastic dynamics. Hence, these networks perform a general form of dynamic sampling tasks based on Bayesian cue integration. While recurrent networks with external stochastic noise (for example [14]) could achieve this computation, the stochastic implementation requires different computational components separately; the probability representation through stochastic inputs and state-dependency of computations via recurrent connections. By contrast, chaotic dynamics unify the two properties through recurrent network interactions in our model.

## Discussion

We demonstrated that the recurrent neural networks trained using the local learning rule can integrate multimodal sensory inputs, compose near Bayes optimal posterior distributions, and draw samples from them by utilizing internally generated chaotic variability. Chaotic dynamics is apparently incompatible with reliable computation because of the butterfly effect that exponentially spreads dynamical trajectories initially started from nearby network states. Therefore, previous works have mostly avoided using a directly chaotic phase in the final state. However, as shown in Figs. 3 d, S4, and S5, this work offers a basis for realizing reliable probabilistic computation using chaotic neural dynamics. We speculate that this mechanism may underlie flexible information processing in the brain accompanying large variability, with the fact that experimental supports for chaotic neural dynamics are already reported [25]. Further evidence during Bayesian computation may become available in the future using advanced experimental techniques such as wide-field imaging [70, 71], dense electrophysiology recordings [72], and neural control by optogenetics [73]; our results suggest that injection of a perturbation to some neurons can drastically change the network dynamics and resulting outputs, but without altering the Bayesian posterior distribution from which the outputs are generated. This prediction could be tested by confirming unaltered task performance despite the sensitivity of network states to perturbations during a statistical task, which is not shared with the noise-based sampling approach [9–12, 14, 74].

We note that the use of chaotic dynamics in recurrent neural networks as a source of random-number generators has been investigated [75, 76]. One work [75] proposes an engineering method, using a hand-crafted network architecture, to solve a static problem without addressing Bayesian cue integration. Another work [76] uses chaotic input generated in a separate network instead of assuming external stochastic input to solve static tasks. Hence, similar to standard generative models, these works assumed networks that learn to process input variability as opposed to generate variability, which seems a less flexible architecture because of separated networks for the representation and random-number generation. Neither of the above works addresses how biological systems could implement such computations, how networks with a generic initial structure can learn the task, or how spontaneous variability is related to evoked variability in the presence of an input. In contrast, we have shown not only that chaotic neural dynamics are useful for stochastic sampling in both static and dynamic tasks but also that generic networks can learn Bayesian computation and sampling through the biologically plausible synaptic plasticity rule. Importantly, as described in Fig. 4, our model suggests that the ensemble of evoked activity during training shapes spontaneous activity, even when the training data do not include the corresponding samples for spontaneous activity. Further, the proposed networks not only express the prior distribution but also approximate conditionally averaged posteriors, such as 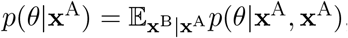, as its generalization. Our results in Fig. 5 showed that the network could represent this distribution for a variety of observed input **x**^A^, simply by omitting input **x**^B^ without needing to explicitly evaluate the computationally intense conditional average. This is an attractive property for brain-inspired computing.

In this work, we suggested the role of chaotic variability in sampling-based computations. However, it might also serve other roles. For example, during training, we applied the node perturbation algorithm that randomly perturbs neural activity and correlates the perturbation with future rewards. While the use of white-noise perturbations is suitable for uniformly exploring the activity space without assumptions, structured chaotic variability, shaped by experience, may be suitable for targeted exploration in the activity space [77]. The use of internally generated chaotic variability to guide the exploration of node perturbation learning and comparison of its performance with the one using white noise is a potential direction for future research. Such structured neural variability might control the trade-off between exploration and exploitation in learning. Notably, while we considered deterministic neural networks without noise for simplicity, chaotic dynamics are well-defined in the presence of noise [66, 78]. We do not rule out a contribution from noise in neural variability; however, we simply focused on the network mechanism that deterministically generates irregular neural dynamics via strong synapses, which could enhance stochastic seeds of variability if any. We believe that future studies will identify their separable contributions and potential synergy. For example, additive noise can control the chaotic transition to maintain high memory capacity [78] and quenched noise can also modulate high-performance regime introducing multistability that includes chaotic dynamics [79].

Several extensions of the proposed model are possible. First, recurrent neural network models have offered interpretable and parsimonious accounts to bridge different types of cognitive behaviors and neural circuit activity [6, 80–83]. Therefore, the Bayesian mechanism proposed with the simple and abstract recurrent neural network model could be tailored to explain more specific behaviors, such as motor control, psychophysics effects, and decision-making [29, 63]. Second, we focused on discrete-time dynamics of neural networks for simplicity; however, extensions toward more biologically plausible models are also possible. For example, continuous-time models share many dynamical properties with our model and may become essential if more precise temporal patterns are needed. Third, we have assumed that neurons communicate with each other through their firing rates, but communication through spiking activity may play an important role. Importantly, the node perturbation learning rule that we used is directly applicable in spiking neural networks as well [54]. It would be interesting to explore whether this difference may contribute to biological correspondence as well as energy and computational efficiency. Fourth, humans and other animals exhibit behavioral variability in their environments. Deterministic models explain the hierarchy of multiple timescales and the interplay among them [84, 85]. Chaotic dynamics may help bridge neural and behavioral variability in the context of probabilistic tasks. Our results suggest that macroscopic variability in animal behaviors may partly arise from probabilistic sampling utilizing microscopic neural activity. Finally, although we only considered five angular positions, higher resolution than 2*π/*5 is realized in the head direction systems in animals (for example [86]), and it does not seem appropriate to scale our computation with a larger readout network using the argmax function. One possible way to address this issue is to increase the number of output neurons but let them interact through fixed lateral couplings to achieve a biological tuning curve [87, 88]. The estimated position may be read out as the center of the activity bump.

## Methods

### Model

A recurrent neural network model, where the *N* neurons are recurrently connected through the synaptic weights denoted as the matrix *J*, and the membrane potential variable of a neuron *i* at time *t* is represented as *h*_*i*_(*t*), was considered. The dynamics of the networks are defined by **h**(*t*): **h**(*t*) = *Jϕ* (**h**(*t −* 1)) + Σ*α*=*A,B K*^*α*^**x**^*α*^ + **c**, where *c*_*i*_ represents the time-independent baseline of the activity of neuron *i* and 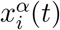 represents the binary state of a sensory neuron *i* of the input population *α*, and fixed input weights are *K*^*α*^ (Figs. 1a and 6 a). The output of the network at time *t* is specified by 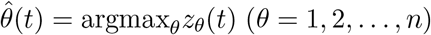 with **z**(*t*) = *Wϕ*(**h**(*t*)) + **b**, where a row of *W*, **W**_*θ*_ is the readout weight for the output *θ*, and *b*_*θ*_ is the readout bias. The activity of the output neurons is thus given by 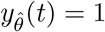 and 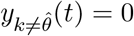.

### Learning rule

We adopted a biologically plausible learning rule, the node perturbation rule [54, 89], which requires only local computation and at the same time approximately reproduces the outcome of the error backpropagation learning. We assume noisy inputs during learning to induce perturbative exploration of synaptic weights and eligibility traces [90]. In this learning rule, the networks exploit the global signals, which can be regarded as “the third factors” [59, 60] and are linked to errors or rewards. Additionally we define the error function as the square of the Hellinger distance between the true posterior distributions and the empirical distributions generated by the network; 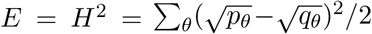 is the Bayesian posterior and *q*_*θ*_ is the generated histogram.

The error was defined as a function of the empirical distribution of network output, which we evaluate as a histogram over a fixed time interval.

During the training period, we perturbed the systems by adding small stochastic noises: the neural dynamics during training are described as

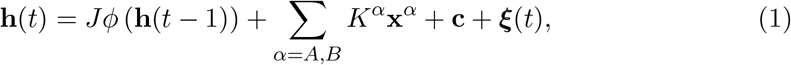

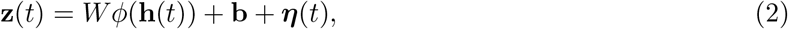

where ***ξ***(*t*) and ***η***(*t*) are independent noises.

As an example, we illustrate the learning of *J*_*ij*_ in detail, while other cases follow similarly. The error function can be expanded over the power of the input trajectory as

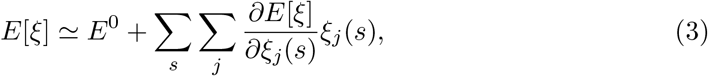

where *E*^0^ denotes the baseline error value without the perturbation input. By averaging the product of *ξ*_*i*_(*t*) with this equation over the statistics of *ξ* with the Dirac delta correlation, we see

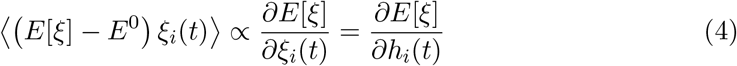

holds. Using this relation, we obtain the expression for the gradient of the error function over *J*_*ij*_:

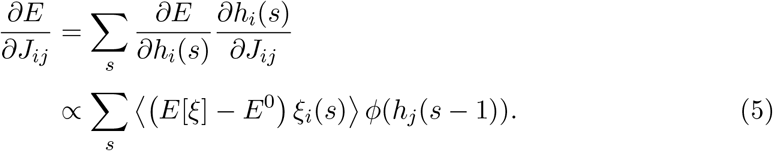

We use the gradient descent method, and the resulting update equation for *J*_*ij*_ is written as

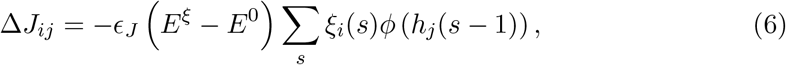

where *E*^*ξ*^ and *E*^0^ represent the error values under the existence and absence of the perturbation, respectively. Here we take the sum over time-step *s* during a batch period. Similarly, for the other adaptive parameters

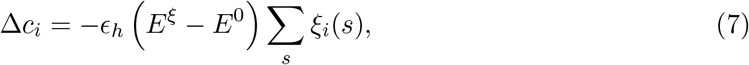

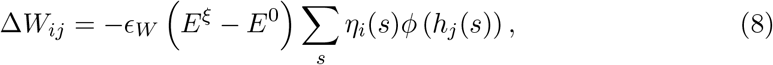

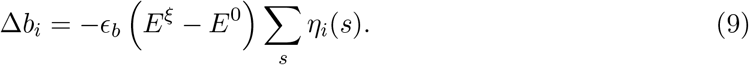

Here denotes the learning rate for each factor. We adopted the Adam method [91] to determine the learning rate, in which the hyperparameters are fixed as usually: ϵ^Adam^ = 0.001, 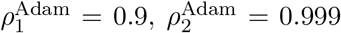, and *δ*^Adam^ = 10^*−*8^ for all parameters *J*, **c**, *W*, and **b**, while other schemes could also be used [45]. Stochastic noises *ξ*_*i*_(*t*) and *ζ*_*i*_(*t*) are drawn from the uniform distribution (for example, in Fig. 2 [*−*1, 1] are their supports). After the training period, the stochastic noises were no longer applied.

### Static cue integration task

To study the capability of the chaotic neural networks in probabilistic computation, we considered a cue integration task where the networks infer the posterior distributions from A and B inputs, as shown in Figs. 1, 2, and 3. The networks received inputs from sensory neurons in populations A and B, whose binary activities (0 or 1) are determined stochastically according to the tuning curves described below.

We considered two different sensory populations A and B, each consisting of *n* = 5 neurons, and the recurrent neural network consisting of *N* = 100 neurons. We set different tuning curves for A and B (see Fig. 1 b): a neuron *i* in population A becomes active with the probability 0.7 for *θ* = *i*, 0.5 for *θ* = *i ±* 1 (mod 5), and 0.3 for *θ* = *i ±* 2 (mod 5), respectively. For example, given the orientation value *θ* = 1, we have the conditional probability of a neuron *i* firing 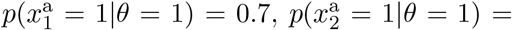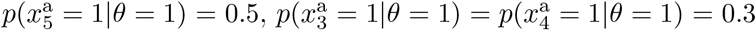. The tuning curve of a neuron *i* in B is specified with the probability 0.8 for *θ* = *i*, 0.5 for *θ* = *i ±* 1 (mod 5), and 0.2 for *θ* = *i ±* 2 (mod 5), respectively. In Figs. 4 and 5, we removed the samples where all neurons in A or B are inactive; if **x**^A*/*B^ = 0 is produced, we draw the **x**^A*/*B^ again. As a result, at least one neuron in A and B would be active and the networks thus receive nonzero stimuli from both populations.

We allocated a transient period of 10 time steps and a generative period of 190 time steps for each input.

In this task, the aim of a network is to generate probabilistic distributions by sampling the output *y*, whose distribution should be as close as possible to the Bayesian posterior of the hidden variable *θ*. After presenting a batch of 50 periods, the error values were calculated, and their means were used to update the weights and biases.

The initial values of internal connectivity *J*_*ij*_ were drawn from a Gaussian distribution with standard deviation 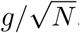, where we set *g* = 8.0 in Figs. 2, 3 a, 4, S2 c-d, and S.3 and *g* = 1.05 unless it is stated otherwise. The readout weights *W*_*ij*_ were initially drawn from a Gaussian distribution with standard deviation 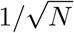. We assumed the zero readout biases *b*_*i*_ = 0 for all network neurons through this task. The input connections *K*_*ij*_ from sensory neurons to the recurrent networks were drawn from a standard normal distribution (10 times larger deviations are used in Fig. S3), and their values were fixed in simulations.

### Temporal statistical task

To study the capacity of the chaotic networks to capture temporal relations by their dynamics, we considered a temporal task in which the recurrent neural networks receive inputs from sensory neurons and generate their outputs so that their sequence follows target transient probabilities.

We considered two groups of sensory neurons A and B. The recurrent neural networks have *N* = 100 neurons as seen in Fig. 6 a. Both sensory neurons take binary states and we trained network dynamics for three of their firing patterns: *x*^A^, *x*^B^ = (0, 0), (1, 0), (0, 1), as spontaneous, A-evoked, B-evoked states, respectively. For each pattern, we set the target transition probabilities that the network should learn, as represented with the red blank boxes in Fig. 6 c below; the target transition probability matrices are denoted concretely as (1*/*3, 1*/*3, 1*/*3; 1*/*3, 1*/*3, 1*/*3; 1*/*3, 1*/*3, 1*/*3) for (*x*^A^, *x*^B^) = (0, 0), (0.25, 0.5, 0.25; 0.25, 0.25, 0.5; 0.5, 0.25, 0.25) for (*x*^A^, *x*^B^) = (1, 0), and (0.3, 0.1, 0.6; 0.6, 0.3, 0.1; 0.1, 0.6, 0.3) for (*x*^A^, *x*^B^) = (0, 1), where the entry of *i*th row and *j*th column denotes the conditional probability *p*(*θ*(*t* + 1) = *i*|*θ*(*t*) = *j*). We trained the network with a time window of 500 time steps for each of the three input patterns.

The gain parameter of synaptic connectivity was set as *g* = 12.5, the initial *K*_*ij*_ was drawn from a standard Gaussian distribution, and initial *W*_*ij*_ was a Gaussian with a standard deviation 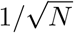. We initially set biases of neurons and readout weights to zero and assumed a nonzero plastic baseline parameter **c** for neural activity.

## Acknowledgements

We thank Johnatan Aljadeff for providing critical comments on the manuscript, which improved the paper significantly. This work was supported by RIKEN Center for Brain Science, the Special Postdoctoral Research Program at RIKEN (Y.T.), Brain/MINDS from AMED under Grant No. JP15dm0207001 (T.T.), MEXT KAKENHI Grant Nos. JP18H05432 (T.T.) and JP19K20365 (Y.T.), and the CRCNS award by the US Department of Energy: DE-SC0022042 (Y.T.).

## Supplementary Information

### Evaluation of statistics of neural sampling

Here we illustrate the convergent property of the neural sampling by the chaotic networks. The trained networks construct probabilistic distributions by sampling the values of the output in time. Fig. S1 shows the dynamics of the network and their Hellinger distances, and the snapshots of the generated distributions at multiple points, where the network and task are the same as Figs. 2 and 3 b and d. The network achieves better performance as the sampling duration increases. The networks require only reasonably short time to obtain statistical approximation for a hidden variable.

**Fig. S1:**
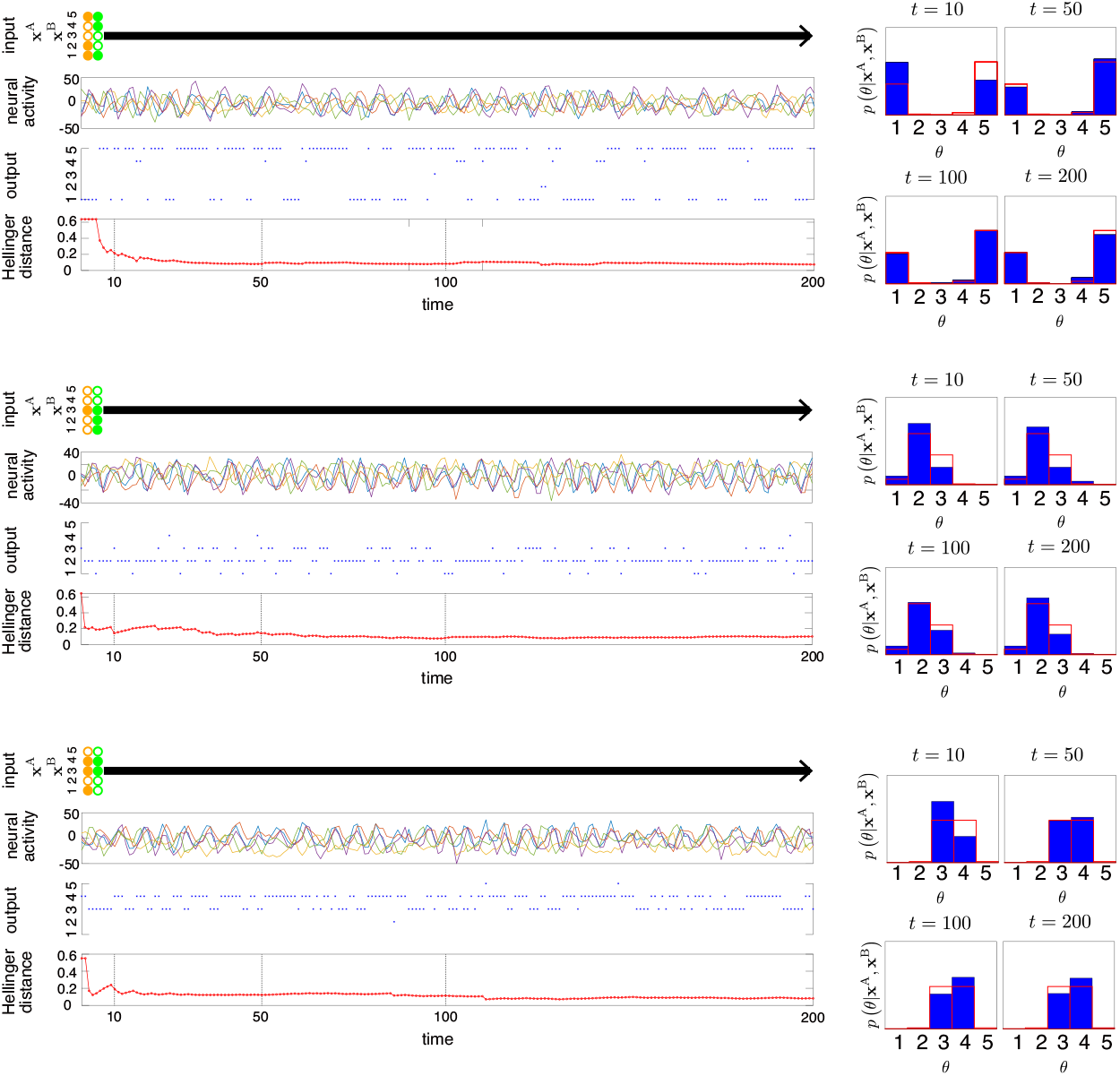
Left: Input, neural activity of some neurons, output, and the Hellinger distance between the output distribution and true distribution in time for 3 examples. Right: Snapshots of the output distribution via neural sampling at *t* = 10, 50, 100, 200, where the red boxes represent the Bayesian posteriors.

### Synaptic changes in internal connectivity through learning

In the main text, we considered the learning scheme in which the synaptic parameters of the networks and readout parameters were trained during the learning period. Here we elaborate on the benefits of synaptic adaptation by showing the case, where only the readout weights were trained as in reservoir computing studies, and compare the result to the one presented in the main text.

We applied the same node perturbation learning rule to the readout weights of the reservoir for two cases, *g* = 1.05 and *g* = 8.0. In the first case (*g* = 1.05), the initial networks are placed on the edge of chaos, in which chaotic dynamics can be suppressed for some input patterns. The other case shows strong chaotic behavior.

The eigenvalues of the weight matrices of the networks before training, which also corresponds to the reservoir computing scheme with fixed *J*, and after training, are shown in Fig. S2, a for *g* = 1.05 and c for *g* = 8.0. The learning curves comparing the proposed scheme with the reservoir computing are shown in Fig. S2, b for *g* = 1.05 and d for *g* = 8.0. When the initial condition is set around the edge of chaos, we see that the learning enhances the chaoticity by significantly changing the internal synaptic weights, which also improved the performance (Fig. S2 a,b). When the initial network lies at the appropriate chaotic phase, the training of internal synaptic weights does not significantly change the eigenvalues, and task performance is similar between the proposed and reservoir computing schemes (Fig. S2 c, d).

However, even when the initial condition is set at *g* = 8.0 as in Fig. S2 c, d, 10 times stronger input weights, which strongly suppress chaos [66], can cause a problem as shown in Fig. S3. In this case, chaos is suppressed for some input patterns, and the training of readout weights alone cannot produce appropriate output distributions. Learning internal synaptic weights with the proposed scheme, however, rescues this failure (Fig. S3). We conclude that training internal synaptic weights adds flexibility even when the initial condition or the network architecture is not fine-tuned.

### Robustness in probabilistic distributions

**Fig. S2:**
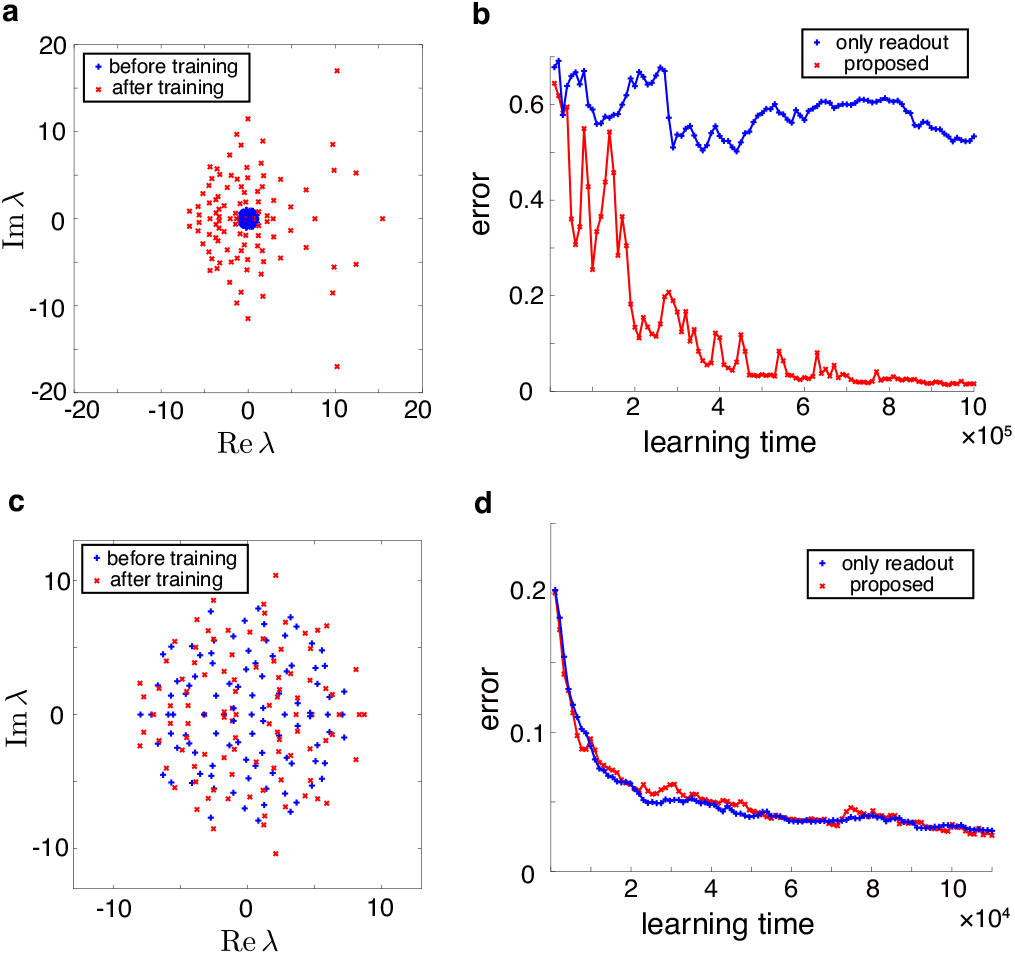
**a, c**, Eigenvalues of coupling matrices before training, which also corresponds to the reservoir computing case with fixed *J*, and after training. **b, d**, Learning curves for the reservoir and proposed schemes. Top: the initial couplings drawn with the gain parameter *g* = 1.05. Bottom: *g* = 8.0.

**Fig. S3:**
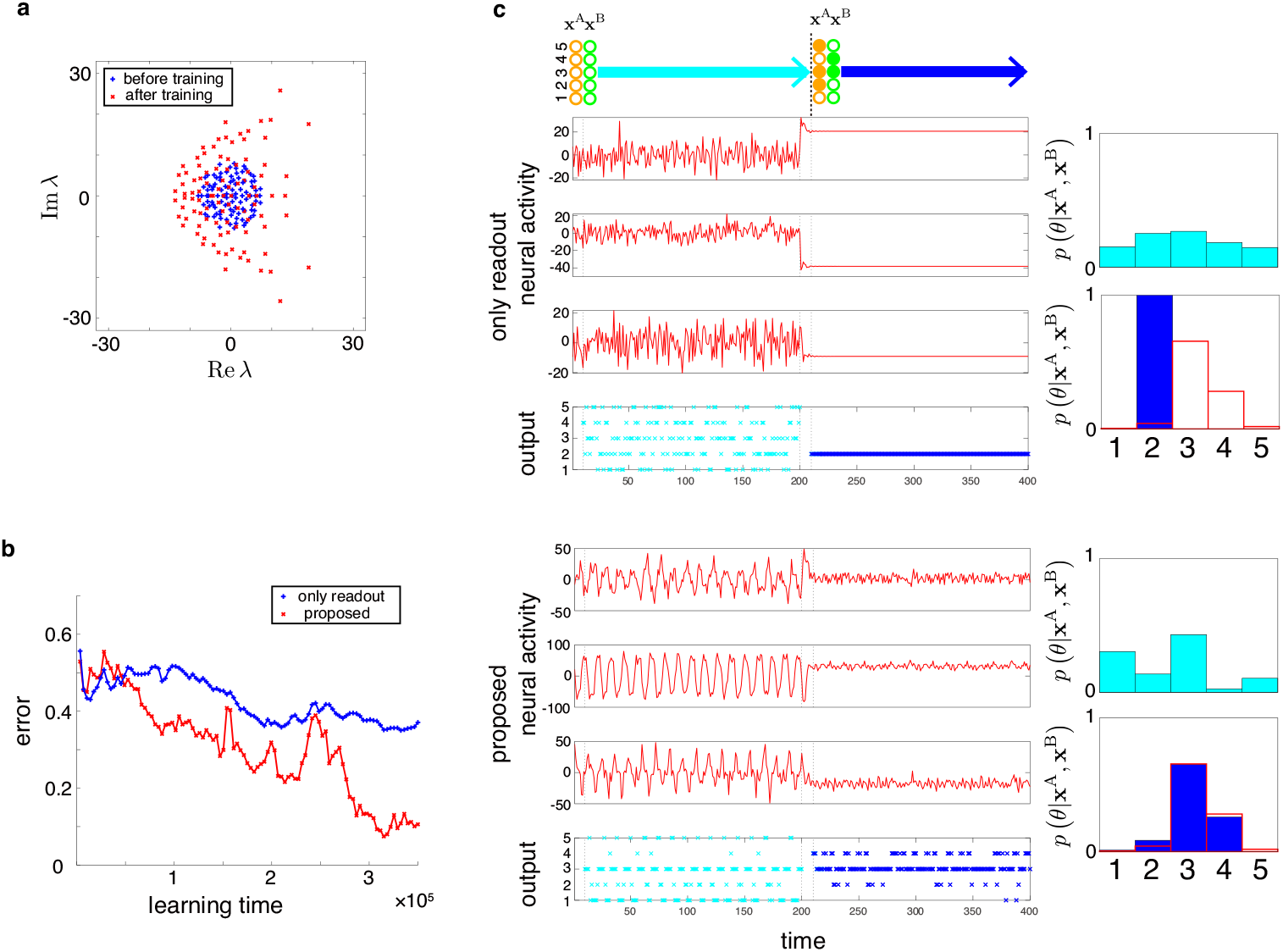
Learning with strong input weights. **a**, Eigenvalues of coupling matrices before training and after training. **b**, Learning curves for the reservoir and proposed schemes. Initial internal synaptic weights were drawn at *g* = 8.0. Ten times stronger input weights than the model in the main text were used. **c**, Input, neural activity, output, and generated distributions in the trained networks over periods without inputs (first half) and with inputs (latter half) for two input patterns, the the reservoir (top) and the proposed cases (bottom) are shown.

As shown in Fig. 2 c, while the chaotic networks showed significantly altered dynamical trajectories in response to activity perturbation, they can still produced similar posteriors. The result highlights the robustness of the proposed probabilistic computation even in the presence of the butterfly effect. Fig. S4 shows the output distributions in response to different sensory input patterns without (blue) and with (magenta) activity perturbation.

**Fig. S4:**
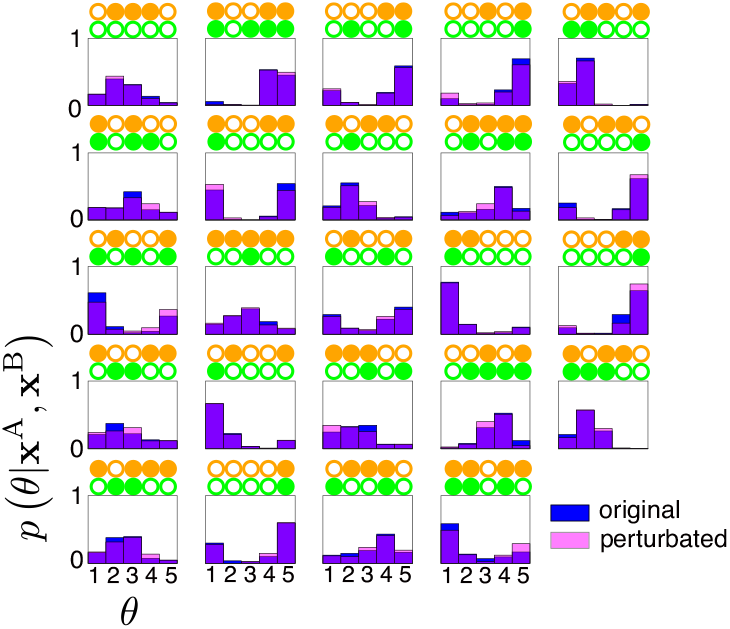
Generated distributions by neural sampling for different sensory input patterns when the network activity is not perturbed (blue) and perturbed (magenta). The trained network is the same as in Fig. 2 b below and Fig. 3 b,d

The above simulation shows robust network performance when the networks exhibit chaotic dynamics characterized by Lyapunov instability, i.e., the sensitivity of network activity to perturbation. We next study the effect of structural perturbation, namely, how a small perturbation to synaptic weights may affect task performance. Figure S5 shows the neural activities, the outputs, and the distributions provided by the original trained and perturbed networks. In contrast, Fig. S5 shows that perturbation to synaptic weights did not affect significantly the network activity and performance. We also observed similar output distributions also in the perturbed case (Fig. S5 b).

**Fig. S5:**
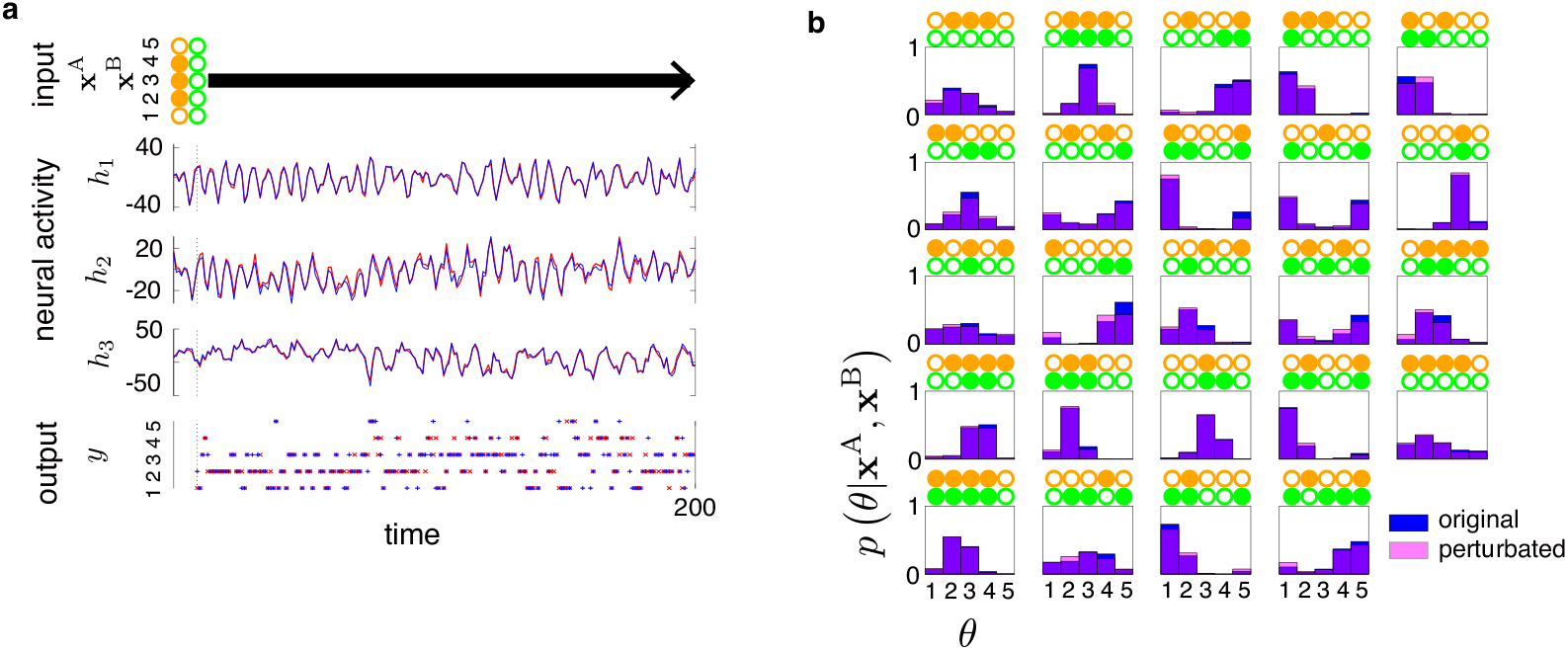
Generated distributions by neural sampling from a trained network that is the same as Fig. 2 b below and Fig. 3 b,d and from the other that has a small perturbation in the synapse parameters. **a**, Input, activity of selected neurons, output in the two networks. **b**, Probabilistic distributions generated by the two networks for several patterns.

